# The transcriptional regulator Lrp activates the expression of genes involved in tilivalline enterotoxin biosynthesis in *Klebsiella oxytoca*

**DOI:** 10.1101/2024.02.29.582825

**Authors:** Miguel A. De la Cruz, Hilda A. Valdez-Salazar, Nayely Robles-Leyva, Tania Siqueiros-Cendón, Quintín Rascón-Cruz, Diana Rodríguez-Valverde, Nancy León-Montes, Jorge Soria-Bustos, Roberto Rosales-Reyes, María L. Cedillo, Jorge A. Yañez-Santos, J. Antonio Ibarra, Javier Torres, Jorge A. Girón, James G. Fox, Miguel A. Ares

## Abstract

The toxigenic *Klebsiella oxytoca* strains secret the tilivalline enterotoxin, which causes antibiotic-associated hemorrhagic colitis. The tilivalline is a non-ribosomal peptide synthesized by enzymes encoded in two divergent operons clustered in a pathogenicity island. The transcriptional regulator Lrp (*l*eucine-responsive *<u>r</u>*egulatory *<u>p</u>*rotein) controls the expression of several bacterial genes involved in virulence. In this work, we determined the transcriptional expression of *aroX* and *npsA*, the first genes of each tilivalline biosynthetic operon in *K. oxytoca* MIT 09-7231 wild-type and its derivatives Δ*lrp* mutant and complemented strains. The results show that Lrp directly activates the transcription of both *aroX* and *npsA* genes by binding to the intergenic regulatory region in a leucine-dependent manner. Furthermore, the lack of Lrp significantly diminished the cytotoxicity of *K. oxytoca* on HeLa cells due to tilivalline reduced production. Altogether, our data highlight Lrp as a new regulator by which cytotoxin-producing *K. oxytoca* strains control the expression of genes involved in the biosynthesis of their main virulence factor.

**IMPORTANCE:** Tilivalline is an enterotoxin that is a hallmark for the cytotoxin-producing *K. oxytoca* strains, which cause antibiotic-associated hemorrhagic colitis. The biosynthesis of tilivalline is driven by enzymes encoded by the *aroX*- and NRPS-operons. In this study, we discovered that the transcriptional regulator Lrp directly activates expression of the *aroX*- and NRPS-operons and, in turn, tilivalline biosynthesis. Our results underscore a molecular mechanism by which tilivalline production by toxigenic *K. oxytoca* strains is regulated and shed further light on developing strategies to prevent the intestinal illness caused by this enteric pathogen.

## INTRODUCTION

*Klebsiella oxytoca* is an opportunistic Gram-negative bacterium that resides in the colon and, after a β-lactam antibiotic treatment, utilizes its constitutively expressed β-lactamase to survive. In contrast, other commensal bacteria are eliminated (1). Pathobiont *K. oxytoca* is the causative agent of antibiotic-associated hemorrhagic colitis (AAHC) (2). Lacking a competitive microbial community, the pathobiont *K*. *oxytoca* originates a dysbiosis (3). Dysbiosis is a condition in which a loss of commensals occurs, leading to an expansion of pathobionts (4, 5). Subsequent expansion of *K*. *oxytoca* in the colon leads to overproduction of tilivalline (TV) cytotoxin. TV is a non-ribosomal peptide that impedes cell cycle progression owing to the augment of nucleation and elongation of tubulin polymerization (6). TV causes apoptosis and damages the tight junction protein claudin-1, causing an impaired intestinal barrier in the colonic epithelium cells (7). *K*. *oxytoca* synthesizes TV via non-ribosomal peptide synthases encoded by the *aroX*- and NRPS-operons, which are part of a pathogenicity island (PAI) (8–10).

Bacteria respond to changes in environmental conditions by regulation of transcription. Indeed, bacteria must rapidly sense and respond to their environment to optimize fitness, adapt, and express their virulence factors in the case of pathogenic bacteria. The global regulator Lrp (*l*eucine-responsive *<u>r</u>*egulatory *<u>p</u>*rotein) is one of the transcription factors that control such responses (11–13). Previously, we found that *aroX*- and NRPS-operons transcription is activated directly by the global regulator CRP (cAMP receptor protein), where the regulation via CRP is boosted by lactose (14) Nevertheless, the role of other global regulators in the expression of *aroX*- and NRPS-operons remains unknown.

Lrp is one of the top seven global regulators involved in the modulation of various metabolic and physiological functions and is widely distributed among prokaryotes and archaea. In *Escherichia coli*, Lrp regulates the expression of approximately one-third of its genome; the overall regulatory behavior of Lrp extends to 38% of *E*. *coli* genes, including some involved in amino acid metabolism and transport (15, 16). Lrp is an 18.8-kDa DNA-binding protein that can act as a positive and negative transcriptional regulator by directly or indirectly binding to the regulatory regions of target genes (16–20). L-leucine can augment its activity; indeed, it has been shown that Lrp self-associates mainly into octamers, and such oligomeric structure could be enhanced by the presence of L-leucine (21–24). Additionally, Lrp monitors a general nutritional state by sensing the concentrations of L-leucine in the cell and regulating genes involved in entering the stationary growth phase in *E. coli* (25, 26).

In this work, we aimed to determine the role of Lrp in the transcriptional regulation of the tilivalline biosynthetic genes in the absence and presence of L-leucine. Besides, we also evaluated the role of Lrp in controlling the cytotoxic effect of *K. oxytoca* on HeLa cells. This study highlights the function of Lrp as a transcriptional activator of genes implicated in the biosynthesis of *K. oxytoca* TV enterotoxin.

## MATERIAL AND METHODS

### Bacterial strains and growth conditions

We used the *K. oxytoca* toxigenic strain MIT 09-7231 (27), its derivative Δ*lrp* mutant and the complemented strain Δ*lrp* pT3-Lrp. All the strains and plasmids used in this work are further listed in Table 1. Tryptone soy broth (TSB) (Difco), and N-minimal medium (N-MM) (5 mM KCl, 7.5 mM (NH_4_)_2_SO_4_, 0.5 mM K_2_SO_4_, 1 mM KH_2_PO_4_, 100 mM Tris-HCl, 10 μM MgCl_2_, 0.5% glycerol, and 0.15% casamino acids) at pH 7.2 (28), were used to grow cultures at 37°C. Culture media were supplemented with 100 μg/mL L-leucine (Sigma) when indicated (29). When necessary, antibiotics were added: ampicillin (200 μg/mL), kanamycin (50 μg/mL), or tetracycline (10 μg/mL).

**TABLE 1.** Bacterial strains and Plasmids used in this study.

| Strains or Plasmid | Description | References |
| --- | --- | --- |
| **Strains** |  |  |
| *K. oxytoca* WT | Wild-type *K. oxytoca* strain MIT 09-7231 | (27) |
| *K. oxytoca* $\Delta lrp$ | *K. oxytoca* $\Delta lrp::Km^R$ | This study |
| *K. oxytoca* $\Delta npsA$ | *K. oxytoca* $\Delta npsA::FRT$ | (14) |
| *E. coli* BL21 (DE3) | F<sup>-</sup> *ompT hsdS<sub>B</sub>* (rB<sup>-</sup>, mB<sup>-</sup>) *gal dcm* (DE3) | Invitrogen |
| *E. coli* MC4100 | Cloning strain<br>F<sup>-</sup> *araD139*Δ(*argF-lac*)<br>*U169 rspL150 relA1 flbB5301 fruA25 deoC1 ptsF25* | (57) |
| **Plasmids** |  |  |
| pMPM-T3 | p15A derivative low-copy number expression vector, *lac* promoter; Tc<sup>R</sup> | (58) |
| pT3-Lrp | pMPM-T3 derivative expressing *lrp* from the *lac* promoter, Tc<sup>R</sup> | This study |
| pMPM-T6 | p15A derivative expression vector, pBAD (*ara*) promoter; Tc<sup>R</sup> | (58) |
| pT6-Lrp | pMPM-T6 derivative expressing N-terminal His<sub>6</sub>-Lrp from pBAD (*ara*) promoter, Tc<sup>R</sup> | This study |
| pKD119 | pINT-ts derivative containing the λ Red recombinase system under | (31) |
|  | an arabinose- inducible promoter, Tc<sup>R</sup> |  |
| pKD4 | pANTsy derivative template plasmid containing the kanamycin cassette for $\lambda$ Red recombination, Ap<sup>R</sup> | (31) |

Km<sup>R</sup>, kanamycin resistance; Tc<sup>R</sup>, tetracycline resistance; Ap<sup>R</sup>, ampicillin resistance.

### Lrp protein sequences analysis

Amino acid sequence alignment of Lrp from *K. oxytoca* MIT 09-7231(GenBank accession number: KMV84644.1) with other selected *Enterobacterales* Lrp homologues was carried out using Clustal Omega (https://www.ebi.ac.uk/Tools/msa/clustalo/) (30).

### Generation of the *lrp* isogenic mutant strain

Deletion of *lrp* gene was performed by the one-step mutagenesis procedure by replacing the target gene with a selectable kanamycin resistance gene marker with the λ-Red recombinase system, as previously described (31). To confirm the authenticity of the mutation, it was characterized by PCR and sequencing.

### Construction of plasmids

Plamids pT3-Lrp and pT6-Lrp were generated to complement the Δ*lrp* mutant strain, and to obtain the His_6_-Lrp recombinant protein, respectively. PCR purified products of *K. oxytoca lrp* gene were obtained with specific primers (Table 2). pT3-Lrp construction was obtained by cloning *lrp* gene into pMPM-T3 vector with XhoI and EcoRI restriction enzymes, whereas pT6-Lrp was generated by cloning *lrp*, containing a nucleotide sequence that encodes a His_6_-tag at N-terminal of Lrp recombinant protein, into pMPM-T6 vector with NcoI and HindIII restriction enzymes. The constructs were verified by DNA sequencing.

**TABLE 2.**
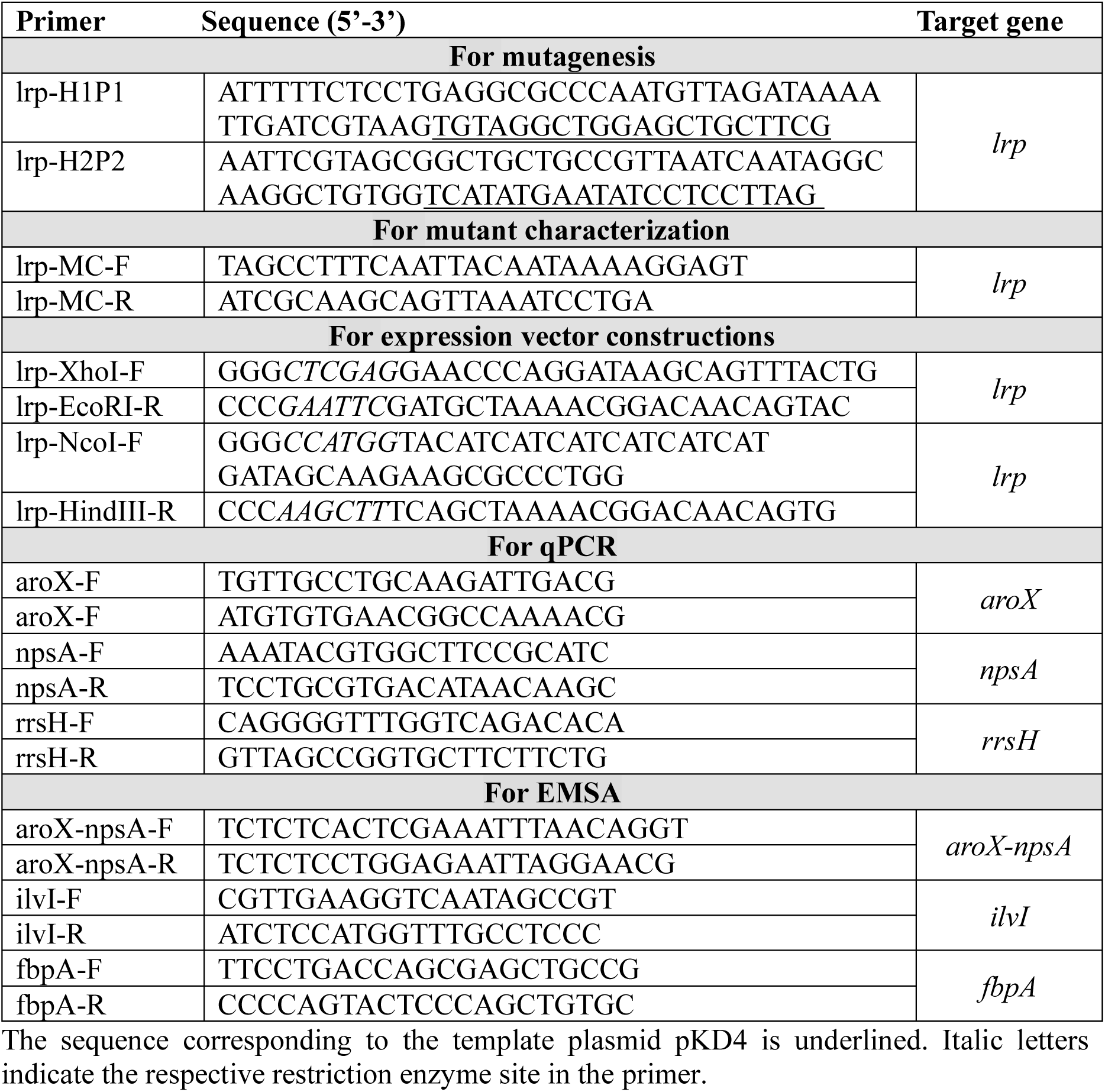
Primers used in this study.

### RNA isolation and Reverse transcription-quantitative PCR

Total RNA was isolated by using the hot phenol method (32). RNA was purified with the TURBO DNA-free kit (Invitrogen), and its concentration and purity were determined using the NanoDrop ONE (Thermo Scientific). RNA integrity was evaluated by electrophoresis in a bleach denaturing 1.5% agarose gel, as described (33). cDNA was synthesized with 1 μg of RNA as template employing the RevertAid First Strand cDNA Synthesis Kit (Thermo Scientific). Quantitative PCR was carried out in a LightCycler 480 instrument (Roche) with specific primers (Table 2) as previously described (14). The *rrsH* gene, which codes for 16S rRNA was used for normalization, and experiments were performed in triplicate from three independent assays. Minus RT controls were included in all experiments. The 2^-DDCt^ formula (34, 35) was used to calculate the relative gene expression.

### Purification of Lrp protein

The His_6_-Lrp expression vector pT6-Lrp was electroporated into competent *E. coli* BL21 (DE3) strain, and the recombinant protein was purified as previously described (36). Briefly, bacteria containing recombinant plasmid was grown to mid-exponential phase; L(+)-arabinose was then added and cultured for 6 h at 37°C. After centrifugation, the cellular pellet was resuspended in urea buffer (8 M urea, 100 mM Na_2_HPO_4_ and 10 mM Tris-HCl, pH 8.0) and sonicated. The lysate was pelleted by centrifugation, and the supernatant was filtered through Ni-NTA agarose column (Qiagen). After washing with 50 mM imidazole (200 mL), the protein was eluted with 500 mM imidazole (10 mL). Analysis of fractions were performed by SDS-PAGE and Coomasie blue staining. Protein concentration was calculated by the Bradford method (Bio-Rad). The recombinant His_6_-Lrp purified protein was stored at -70°C.

### Recognition of promoters and Lrp binding sites on the *aroX*-*npsA* intergenic region

The prediction of promoters on the *aroX* and *npsA* regulatory regions were determined by analyzing 430 nucleotides upstream to the initial codon of both genes by using the genomic sequence from *Klebsiella oxytoca* MIT 09-7231 (GenBank accession number: GCA_001078175.1) with the web-based program *Neural Network Promoter Prediction* (https://www.fruitfly.org/seq_tools/promoter.html). The Lrp putative motifs were identified by using the web-based softwares *Transcription Factor Binding Site Prediction* (TFBS) (http://genome2d.molgenrug.nl/g2d_pepper_TFBS.php), and *Softberry BPROM* (http://www.softberry.com/berry.phtml?topic=bprom&group=programs&subgroup=gfindb)

### Electrophoretic Mobility Shift Assays (EMSA)

EMSA experiments were carried out as previously described (14). 100 ng of a 448-bp DNA probe corresponding to the intergenic regulatory region of the divergent *aroX* and *npsA* genes was mixed with increasing concentrations of His_6_-Lrp with 7.0 mM L-leucine (37). As positive and negative controls, DNA probes from regulatory region of *K. oxytoca ilvI* gene and coding region of *M. tuberculosis fbpA* gene, were used. Reactions were incubated at room temperature for 20 min and then separated by electrophoresis in 6% non-denaturing polyacrylamide gels using 0.5X Tris-borate-EDTA buffer. The DNA bands were stained with ethidium bromide and visualized under UV light.

### LDH cytotoxicity assays

The LDH Cytotoxicity Assay Kit (Invitrogen) was used according to the manufactureŕs instructions to measure Lactate dehydrogenase (LDH) released from HeLa cells after damage of plasma membrane integrity. 10 µL of negative control (PBS), culture medium control (TLB), positive control (lysis buffer), and filtered bacterial supernatants (wild-type, Δ*lrp*, Δ*lrp* pT3-Lrp, and Δ*npsA*), were added to 1×10^4^ HeLa cells previously cultivated in 100 µl DMEM high glucose (4.5 g/l) (Invitrogen) with 10% FBS (Gibco) into 96-well flat-bottom plate culture. Then, it was incubated at 37°C under a 5% CO_2_ atmosphere for 48 h. Afterwards, 50 µL each sample medium was transferred to a new 96-well plate, and kit solutions added into each well. The absorbance was measured at 490 nm and 680 nm with a spectrophotometer (Multiskan Ascent, Thermo Scientific). The experiment was performed in three independent biological replicates by triplicates, and results were expressed as LDH cytotoxicity by deducting the 680 nm absorbance background value from the 490 nm absorbance value.

### Statistical analysis

For statistical differences, unpaired two-tailed Student’s *t*-test were performed with the GraphPad Prism 9.0 software (GraphPad Inc., San Diego, CA, United States). All data were obtained from three independent experiments performed by triplicate, and values of *p*<0.005 were considered significant.

## RESULTS

### Lrp amino acid sequence analysis

We initiated this study by comparing the amino acid sequences of a group of *Enterobacterales* Lrp proteins with the *K. oxytoca* toxigenic strain MIT 09-7231 Lrp protein. *K. oxytoca* Lrp amino acid sequence shared an average of 98.24% identity with the other Lrp sequences of *K. pneumoniae, E. coli, Enterobacter cloacae, Salmonella* Typhi, *Shigella dysenteriae, Yersinia enterocolitica, Proteus mirabilis*, and *Serratia marscesens* (Figure 1). The putative helix-turn-helix (HTH) and the βαββαβ-fold motifs were identified at the N- and C-terminal of K. oxytoca Lrp protein, respectively, through comparison with *E. coli* Lrp (38) by using the web-based PROSITE bioinformatic tool (http://prosite.expasy.org). When comparing the amino acid sequences of Lrp from *K. oxytoca* and *E. coli*, 99.39% identity and 100% similarity were obtained, since there is only one change in residue 95 of the protein, being serine for *K. oxytoca* Lrp and threonine for *E. coli* Lrp, both amino acids with uncharged polar side chains containing aliphatic hydroxyl groups.

**Figure 1.**
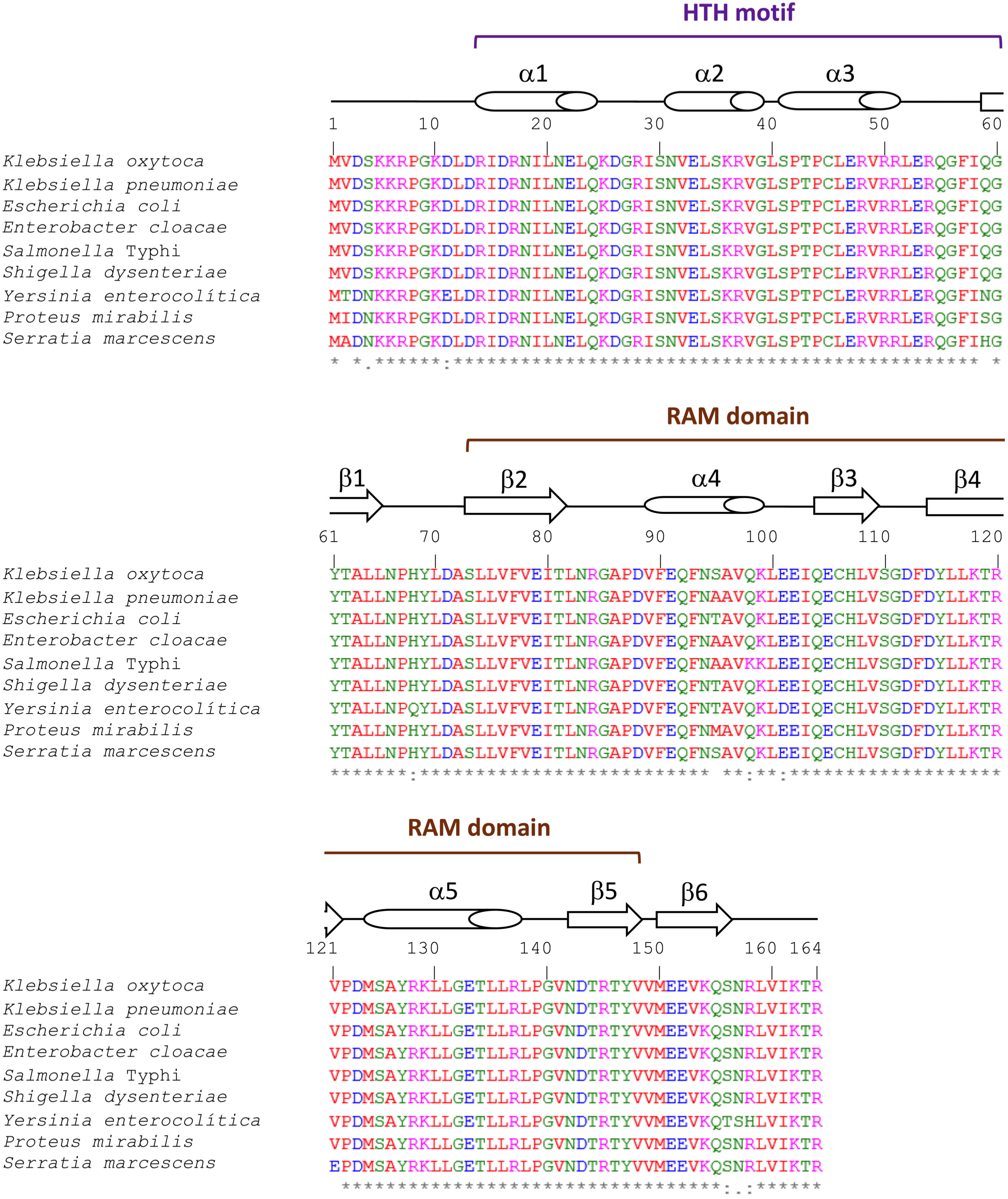
Amino acid sequence alignment of Lrp from *K. oxytoca* and other *Enterobacterales*. The amino acid sequence of Lrp (KMV84644.1) from *K. oxytoca* MIT 09-7231, and Lrp homologs proteins: (BAH62617.1) from *K. pneumoniae* NTUH-K2044, (NP_415409.1) from *E. coli* K-12 MG1655, (ADF62302.1) from *Enterobacter cloacae* ATCC 13047, (AA069588.1) from *Salmonella enterica* serovar Typhi Ty2, (ABB62447.1) from *Shigella dysenteriae* Sd197, (CAL11603.1) from *Yersinia enterocolitica* 8081, (CAR41630.1) from *Proteus mirabilis* HI4320, and (CDG11553.1) from *Serratia marcescens* Db11 were aligned by using Clustal Omega (https://www.ebi.ac.uk/Tools/msa/clustalo/). The predicted regions to encode the HTH motif (DNA-binding), and RAM domain (Regulation of amino acid metabolism) are indicated. Secondary structural elements are shown by barrels (α-helix) and arrows (β-sheet). Hydrophobic, polar, positively charged, and negatively charged amino acids are represented in red, green, magenta, and blue, respectively. The asterisk (*), colon (:), and dot (.) indicate identical, conserved, and semi-conserved amino acids among all aligned sequences.

### Lrp is required for optimal growth of *K. oxytoca* under nutrient limiting conditions

To find out whether *K. oxytoca* toxigenic strain MIT 09-7231 Lrp participates in regulatory process related to growth, we compared growth rates in the wild-type, its derivative Δ*lrp* mutant, and complemented strains in two different culture media: TSB (nutrient rich) and N-MM (nutrient limiting) in the absence and presence of 100 µg/mL L-leucine (29). There were no growth differences between the wild-type, Δ*lrp*, and complemented strains in TSB without or with leucine (Figures 2A, B). In contrast, in N-MM without leucine the Δ*lrp* strain was attenuated in exponential and stationary phases (Figure 2C). The addition of leucine to N-MM restored the growth defect in stationary phase (Figure 2D).

**Figure 2.**
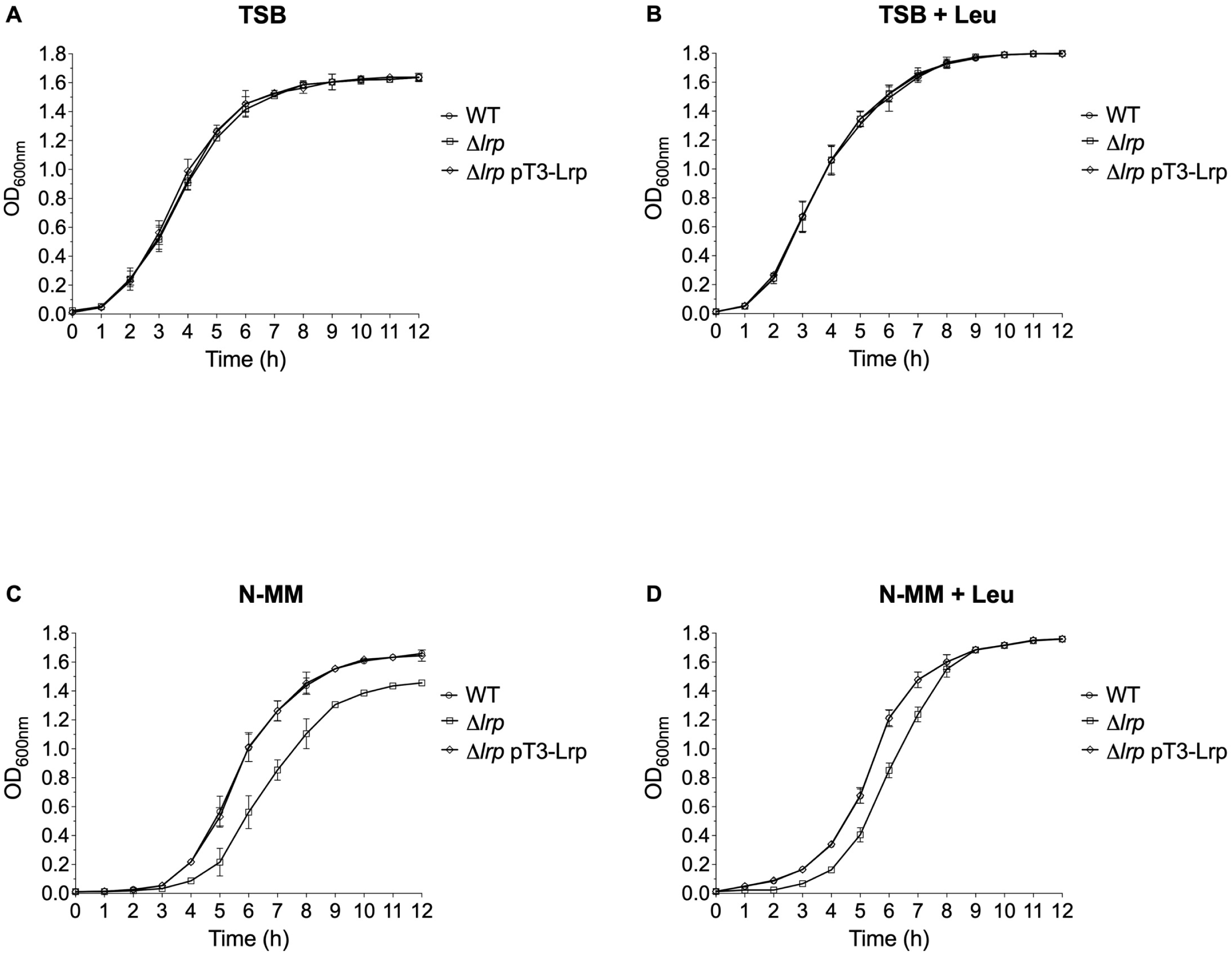
Growth curves of *K. oxytoca* wild-type (WT), Δ*lrp* mutant and Δ*lrp* pT3-Lrp complemented strains. Cultures were grown for 12 h in TSB (nutrient-rich) (**A**, **B**), and N-MM (nutrient-limiting) (**C**, **D**) medium, in the absence and presence of L-leucine (Leu). The OD_600nm_ values were recorded every hour. Data represent the mean of three independent experiments with standard deviations.

These observations indicate that Lrp is involved in *K. oxytoca* toxigenic strain MIT 09-7231 growth regulatory processes as in other bacteria (39–42). Furthermore, the addition of leucine improved growth suggesting that this amino acid may play an important role in maintenance of central metabolism in *K. oxytoca* as has been reported in another microorganism (43–45).

### Lrp is an activator of *aroX* and *npsA* genes transcription

To investigate the regulatory activity of Lrp on *aroX* and *npsA* transcription, expression of such TV genes was determined by RT-qPCR in *K. oxytoca* toxigenic strain MIT 09-7231. In TSB without leucine the transcriptional levels of both *aroX* and *npsA* genes were down-regulated 3-fold and 2-fold, respectively, in the Δ*lrp* strain as compared to the wild-type (Figure 3A). The addition of leucine to TSB enhanced the transcription of *aroX* and *npsA* 3.3- and 2.5-fold, respectively, in the wild-type strain. Interestingly, the transcriptional levels of both TV genes were not significantly different in the Δ*lrp* strain either in the absence or the presence of leucine (Figure 3A).

**Figure 3.**
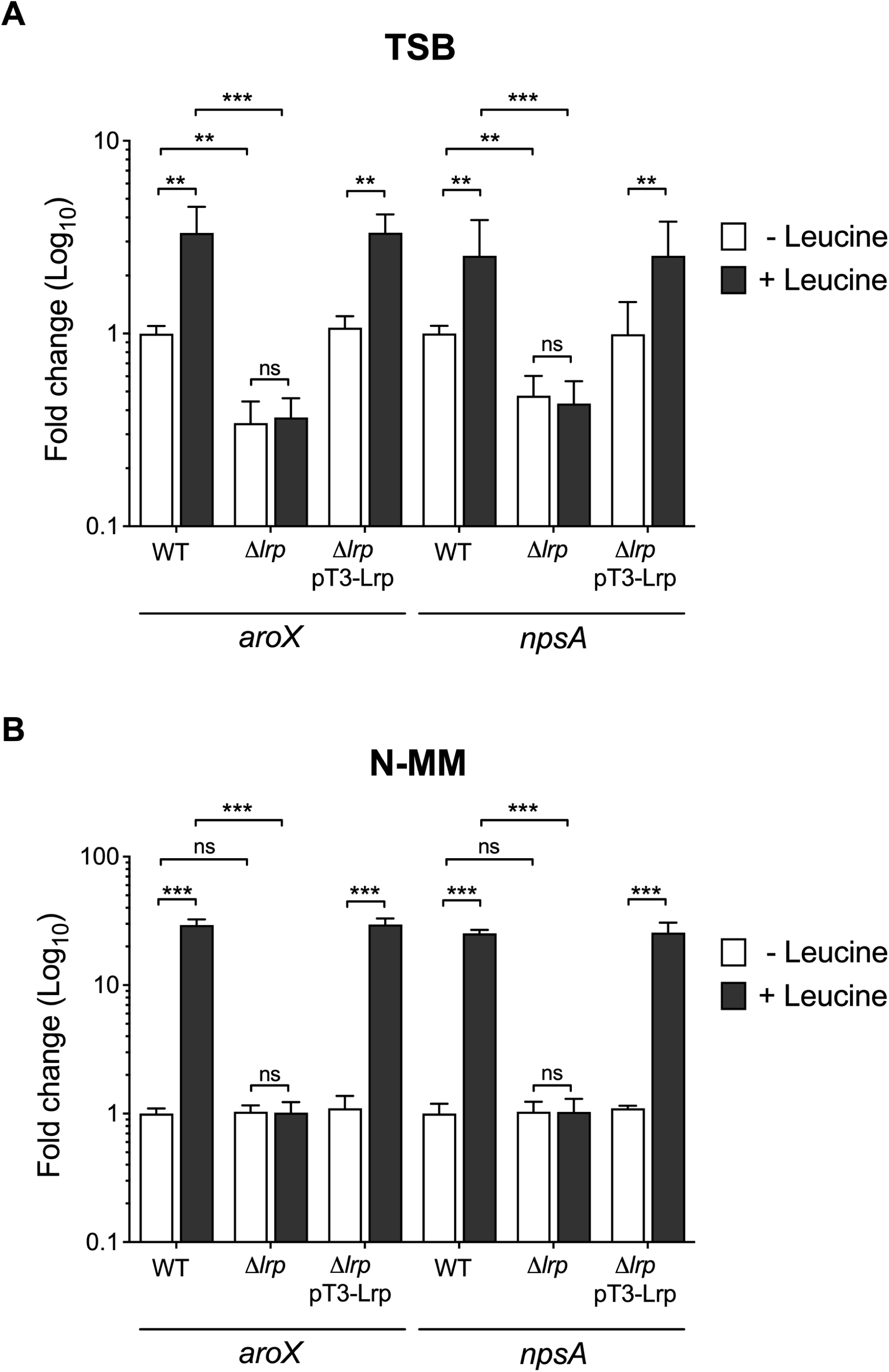
Transcriptional regulation of Lrp on *aroX* and *npsA* genes. Determination of gene expression by RT-qPCR of *aroX* and *npsA* of *K. oxytoca* wild-type (WT), Δ*lrp* mutant and Δ*lrp* pT3-Lrp complemented strains grown in TSB (nutrient-rich) and N-MM (nutrient-limiting) medium at stationary phase (OD_600nm_=1.6) at 37°C, in the absence and presence of L-leucine. Data represent the mean of three independent experiments performed in triplicate with standard deviations. Statistically significant: \*\**p*<0.01; \*\*\**p*<0.001; ns: not significant. All *p* values were determined using unpaired two-tailed Student’s *t*-test.

In N-MM without leucine the transcription of *aroX* and *npsA* was not affected in the Δ*lrp* strain with respect to the wild-type strain. Nevertheless, when leucine was added to N-MM the transcription of both TV genes was up-regulated more than 25-fold in the wild-type strain (Figure 3B). As in TSB, the transcription of *aroX* and *npsA* was not altered in the Δ*lrp* strain in presence of leucine (Figure 3B). In both culture media, TSB and N-MM, the transcription of *aroX* and *npsA* was boosted in the wild-type strain after the addition of leucine, the transcription of both TV genes did not change in the Δ*lrp* strain in the presence of leucine, and the positive regulatory activity of Lrp was only observed in N-MM with leucine (Figures 3A, B). In all cases, transcription of complemented strain harboring the pT3-Lrp plasmid was restored to wild-type levels. These data indicate that Lrp positively regulates the transcription of *aroX* and *npsA* genes. Furthermore, leucine induces Lrp-mediated *aroX* and *npsA* genes transcription.

### Identification of putative promoters and Lrp binding motifs on the *aroX*-*npsA* intergenic region

The cytotoxin biosynthetic gene cluster is arranged in two divergent operons, *aroX* and NRPS (Figure 4A). The *aroX* operon is constituted by the *aroX*, *dhbX*, *icmX*, *adsX*, and *hmoX* genes, whereas the N RPS operon is constituted by the *npsA*, *thdA*, and *npsB* genes (8, 9, 14). To identify regulatory motifs in the intergenic region of *aroX* and *npsA* genes, *in silico* analyses were performed. We identified putative promoters for the *aroX* and *npsA* genes located 151 pb, and 163 pb upstream from their coding regions, respectively (Figures 4B,C). Moreover, two putative Lrp binding motifs were found for both TV operons. The Lrp binding motif for the *aroX* gene was located 105 pb upstream of the putative transcriptions start site (TSS, +1) (Figure 4B), and for the *npsA* gene it was located 88 pb upstream of the TSS (Figure 4C).

**Figure 4.**
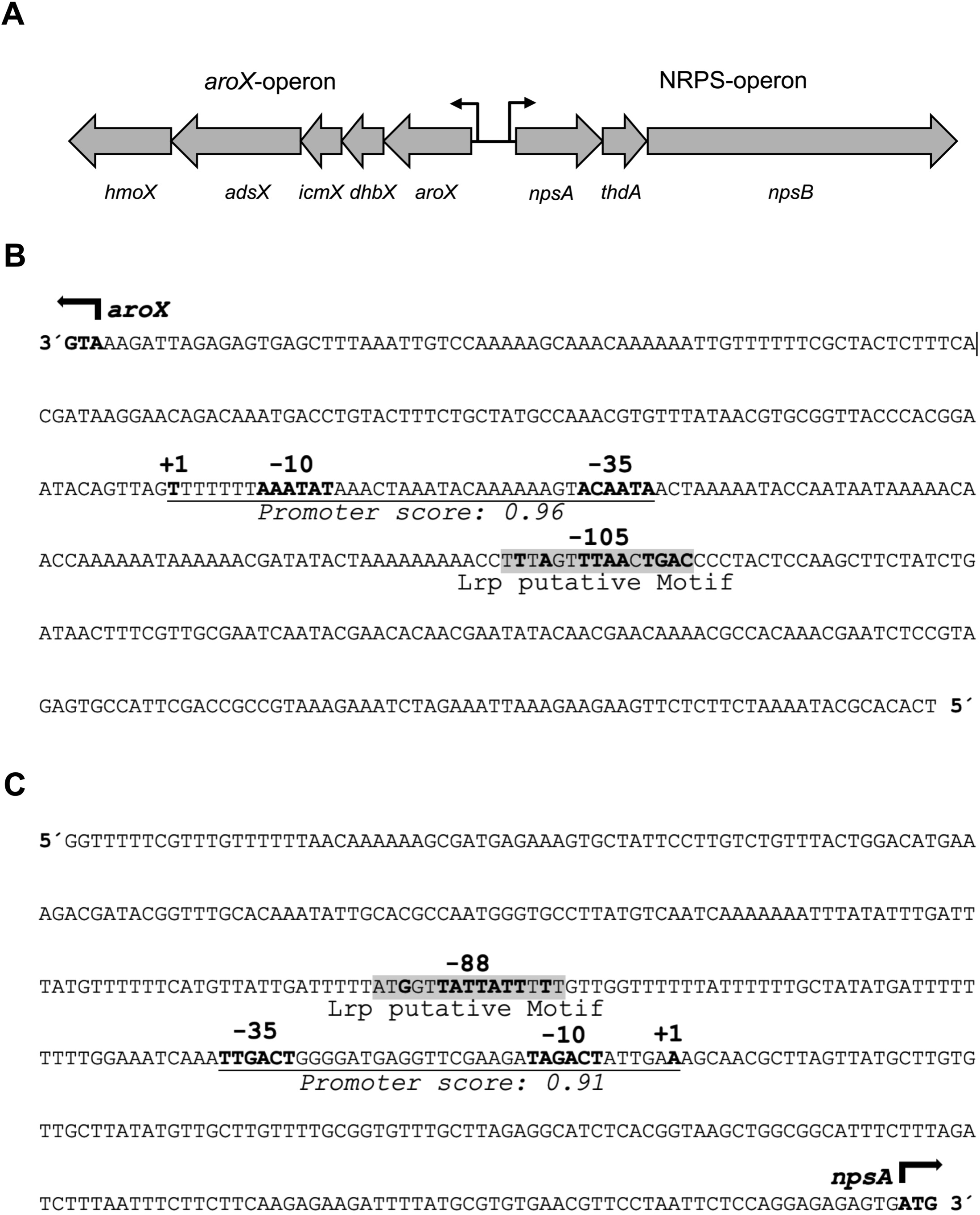
*In silico* analysis of the intergenic region of *aroX* and *npsA* genes. (**A**) Genetic organization of the *aroX* and NRPS operons. Regulatory regions of *aroX* (**B**), and *npsA* (**C**) indicating the initiaton codon (ATG, bold), the predicted promoter region (-35 and -10, bold), the transcription start site (+1, bold), and the Lrp-binding motif (highlighted in grey; nucleotides matching the consensus sequence from *E. coli* are bold).

### Lrp binds to the *aroX*-*npsA* intergenic regulatory region

EMSA were carried out with a recombinant His_6_-Lrp protein and several DNA fragments. Lrp bound to the fragment corresponding to *aroX*-*npsA* intergenic region only in the presence of L-leucine (7 mM) by using 0.4 and 0.8 μM of the recombinant protein (Figures 5A, B). As positive and negative controls, DNA fragments of *ilvI* regulatory region (46, 47) and *fbpA* coding region (48) were employed (Figures 5C, D).

**Figure 5.**
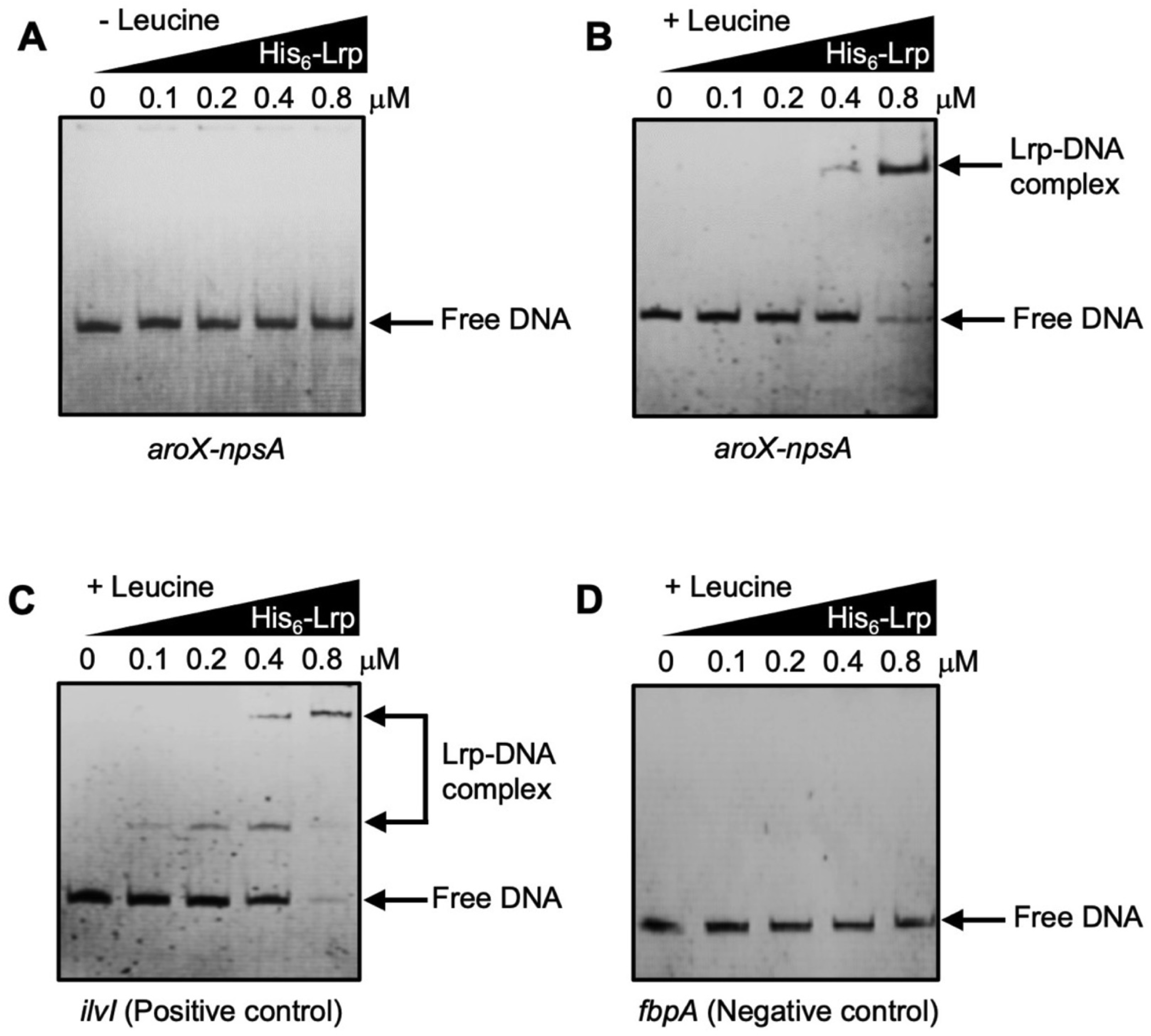
Lrp binds to the intergenic region of *aroX* and *npsA* genes in the presence of leucine. Electrophoretic mobility shift assays (EMSA) were carried out to find out the binding of the His_6_-Lrp purified recombinant protein to the DNA probe from the intergenic regulatory region of *aroX* and *npsA*, in the absence (**A**) or presence of leucine (**B**). DNA probes from *K. oxytoca ilvI* regulatory region (**C**), and *M. tuberculosis fbpA* coding region (**D**) were employed as positive and negative controls, respectively. 100 ng of DNA fragments were individually mixed and incubated with increasing concentrations of purified His_6_-Lrp. L-leucine was added at a final concentration of 7 mM. Arrows show free DNA or Lrp-DNA complex stained with ethidium bromide.

These observations demonstrate that transcriptional activation of *aroX* and *npsA* genes is directly by the bound of Lrp to the *aroX*-*npsA* intergenic region, and such positive regulatory activity is dependent of leucine.

### Lrp promotes the cytotoxic effect of *K. oxytoca* on epithelial cells

To ascertain the regulatory activity of Lrp on the *aroX* and *npsA* transcription, the TV-mediated cytotoxicity was analyzed by the LDH release activity assay on HeLa cells using supernatants from *K. oxytoca* toxigenic strain MIT 09-7231 (wild-type), Δ*lrp* isogenic mutant, and complemented Δ*lrp* strain. The wild-type strain supernatants caused death of HeLa cells, and the cytotoxicity was higher when bacteria were cultivated in media with leucine (Figure 6). In contrast, when supernatants of the Δ*lrp* strain were used, the cytotoxicity was remarkably diminished when bacteria were cultivated in TSB either in the absence or the presence of leucine (Figure 6A). Interestingly, when bacteria were cultivated in N-MM the cytotoxicity only was significantly reduced in the presence of leucine as compared to wild-type strain (Figure 6B).

**Figure 6.**
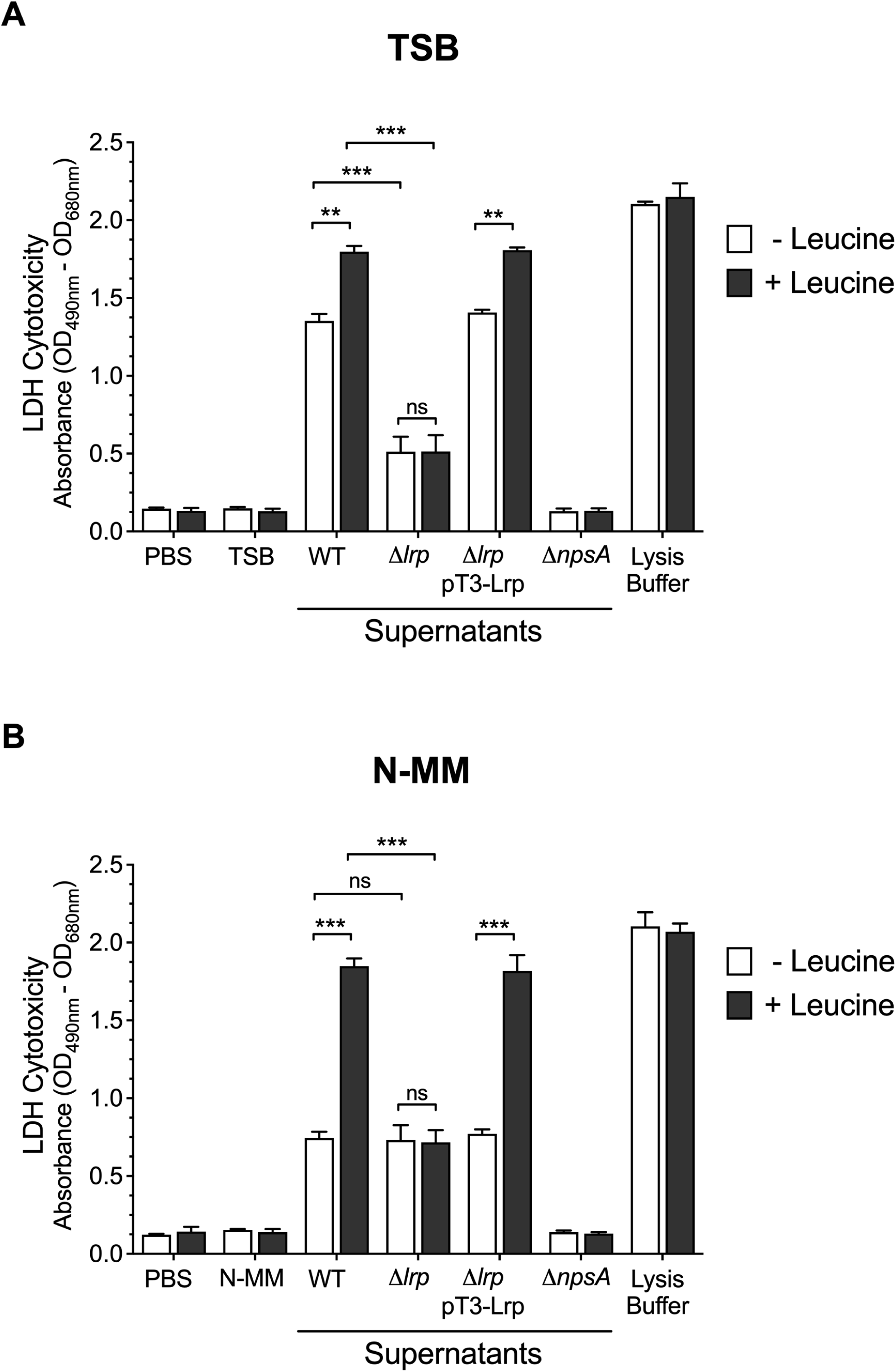
LDH Cytotoxicity of *K. oxytoca* WT, Δ*lrp* mutant, and Δ*lrp* pT3-Lrp strains. HeLa cells were inoculated with TSB and N-MM culture media, and with *K. oxytoca* supernatants (WT, Δ*lrp*, Δ*lrp* pT3-Lrp, and Δ*npsA*), in absence and presence of leucine, for 48 h. After treatment, measurement of extracellular LDH was quantified. Minimal and maximal measurable LDH release was determined by incubating HeLa cells with PBS (negative control), and lysis buffer (positive control), respectively. Statistically significant: \*\**p* < 0.01; \*\*\**p* < 0.001; ns: not significant. All *p* values were determined using unpaired two-tailed Student’s *t*-test.

The cytotoxicity was not affected by the presence of leucine in the Δ*lrp* strain, and the complemented strain supernatants produced LDH release very similar to those of wild-type strain supernatants. Neither culture media nor the non-toxigenic Δ*npsA* strain supernatants produced LDH release (Figures 6A, B). These phenotypic data demonstrate that the Lrp transcriptional regulator activates the expression of TV genes and, consequently, the biosynthesis of TV in *K. oxytoca*.

## DISCUSSION

The global transcriptional regulator Lrp is a member of the feast/famine regulatory proteins, is present in Gram-negative, Gram-positive and archaea, and its amino acid sequence is highly conserved in enteric microorganisms (38, 42, 46, 49). In this study, we found that *K. oxytoca* Lrp protein is homologous to other Lrp proteins from several *Enterobacterales* such as *K. pneumoniae*, *E. coli*, *Enterobacter*, *Salmonella*, *Shigella*, *Yersinia*, *Proteus* and *Serratia*. In terms of bacterial fitness, the lack of Lrp did not affect the growth of *K. oxytoca* in a rich medium (TSB). Nevertheless, in minimal medium (N-MM) the growth of the Δ*lrp* mutant strain was delayed in both exponential and stationary phases. Interestingly, the addition of leucine to the culture medium restored the growth of the Δ*lrp* mutant strain in stationary phase, which agrees with previous reports in *E. coli*, where the growth of a *lrp* mutant strain in minimal medium was also affected with respect to wild-type strains, and such growth defect was restored after addition of leucine (39, 40). Our findings suggest that Lrp in *K. oxytoca*, as in other microorganisms, integrates nutritional signals by regulating the expression of genes required for growth under conditions of amino acid starvation (16, 42, 50, 51).

Regarding transcriptional regulation, Lrp acts either as an activator or as a repressor and mediates a global response to leucine (51). In this work, we determined the transcription of *aroX* and *npsA* genes, which are the first genes of the *aroX* and NRPS operons, respectively. In such operons are encoded the enzymes that participate in the biosynthesis of TV (8, 9). Our results demonstrated that Lrp activates the transcription of *aroX* and *npsA* genes and such regulatory activity is dependent on the presence of leucine as previously reported for other genes such as *oppA*, *sdaA*, *tdh, foo* an*d livK* in *E. coli* (19, 23, 50), and *invF* in *S. enterica* serovar Typhimurium (52).

It has been reported that for some genes, leucine binding to Lrp may increase its affinity to promoters for some genes, leucine binding to Lrp may increaseits affinity to promoters on DNA and, consequently, regulates their expression (Ziegler and Freddolino, 2021). In fact, the binding of leucine to Lrp, not bound to DNA, induces conformational changes, favoring the dissociation of a hexadecameric to octameric form by enhancing dimer-dimer interactions and promoting its binding to DNA (Chen et al., 2005). In our results, the addition of leucine to Lrp recombinant protein was crucial for its DNA-binding activity to the *aroX*-*npsA* intergenic region, suggesting a concerted mechanism of Lrp regulatory protein as was observed with the *fimAICDFGH* promoter (20, 24) Moreover, the position of Lrp-binding sites located -46 and -75 bp of the putative transcription start sites of *aroX* and *npsA* genes, respectively, suggests that Lrp would act as a classic transcriptional activator recruiting to RNA polymerase.

The main virulence determinant of toxigenic *K. oxytoca* is the tilivalline enterotoxin, which causes cytotoxicity on epithelial cells (9, 10, 14, 27, 53–55)The absence of Lrp regulatory protein affected the transcription and subsequently the secretion of tilivalline, resulting in the lack of citotoxicity of *K. oxytoca* toxigenic strain MIT 09-7231. In terms of genes encoding toxins, Lrp has been described as a positive and negative regulator of *xhlA* and *tcdAB* genes, which encodes XhlA hemolysin in *Xenorhabdus nematophila* and toxins A/B in *Clostridium difficile*, respectively (42, 56).

This study describe to Lrp as a novel positive regulator for the expression and production of tilivalline. Current studies are focused on the spatiotemporal regulation of tilivalline by other global and local regulators in the context of pathogenicity of toxigenic *K. oxytoca*.

## AUTHOR CONTRIBUTIONS

MAD, HAV-S, and MAA conceived and designed the experiments. MAD, HAV-S, NR-L, DR-V, NL-M, and RR-R performed the experiments. MAD, HAV-S, TS-C, QR-C, JS-B, MLC, JAY-S, JAI, JT, JAG, JGF, and MAA analyzed the data. MAD, HAV-S, and MAA wrote the manuscript. All authors contributed to the article and approved the submitted version.

## ACKNOWLEDGMENTS

NR-L was supported by a fellowship from CONAHCYT (778751) to complete the Masteŕs degree in Biotechnology from Universidad Autónoma de Chihuahua.

## REFERENCES

1. Alexander EM, Kreitler DF, Guidolin V, Hurben AK, Drake E, Villalta PW, Balbo S, Gulick AM, Aldrich CC. 2020. Biosynthesis, Mechanism of Action, and Inhibition of the Enterotoxin Tilimycin Produced by the Opportunistic Pathogen *Klebsiella oxytoca*. ACS Infect Dis 6:1976–1997.

2. Högenauer C, Langner C, Beubler E, Lippe IT, Schicho R, Gorkiewicz G, Krause R, Gerstgrasser N, Krejs GJ, Hinterleitner TA. 2006. *Klebsiella oxytoca* as a Causative Organism of Antibiotic-Associated Hemorrhagic Colitis. New England Journal of Medicine 355:2418–2426.

3. Herzog KAT, Schneditz G, Leitner E, Feierl G, Hoffmann KM, Zollner-Schwetz I, Krause R, Gorkiewicz G, Zechner EL, Hogenauer C. 2014. Genotypes of *Klebsiella oxytoca* Isolates from Patients with Nosocomial Pneumonia Are Distinct from Those of Isolates from Patients with Antibiotic-Associated Hemorrhagic Colitis. J Clin Microbiol 52:1607–1616.

4. Maharshak N, Packey CD, Ellermann M, Manick S, Siddle JP, Huh EY, Plevy S, Sartor RB, Carroll IM. 2013. Altered enteric microbiota ecology in interleukin 10-deficient mice during development and progression of intestinal inflammation. Gut Microbes 4:316–324.

5. Walker WA. 2017. Dysbiosis, p. 227–232. In The microbiota in gastrointestinal pathophysiology. Elsevier.

6. Unterhauser K, Pöltl L, Schneditz G, Kienesberger S, Glabonjat RA, Kitsera M, Pletz J, Josa-Prado F, Dornisch E, Lembacher-Fadum C, Roier S, Gorkiewicz G, Lucena D, Barasoain I, Kroutil W, Wiedner M, Loizou JI, Breinbauer R, Díaz JF, Schild S, Högenauer C, Zechner EL. 2019. *Klebsiella oxytoca* enterotoxins tilimycin and tilivalline have distinct host DNA-damaging and microtubule-stabilizing activities. Proceedings of the National Academy of Sciences 116:3774–3783.

7. Hering, Fromm, Bücker, Gorkiewicz, Zechner, Högenauer, Fromm, Schulzke, Troeger. 2019. Tilivalline- and Tilimycin-Independent Effects of *Klebsiella oxytoca* on Tight Junction-Mediated Intestinal Barrier Impairment. Int J Mol Sci 20:5595.

8. Dornisch E, Pletz J, Glabonjat RA, Martin F, Lembacher-Fadum C, Neger M, Högenauer C, Francesconi K, Kroutil W, Zangger K, Breinbauer R, Zechner EL. 2017. Biosynthesis of the Enterotoxic Pyrrolobenzodiazepine Natural Product Tilivalline. Angew Chem Int Ed Engl 56:14753–14757.

9. Schneditz G, Rentner J, Roier S, Pletz J, Herzog KAT, Bücker R, Troeger H, Schild S, Weber H, Breinbauer R, Gorkiewicz G, Högenauer C, Zechner EL. 2014. Enterotoxicity of a nonribosomal peptide causes antibiotic-associated colitis. Proc Natl Acad Sci U S A 111:13181–13186.

10. Tse H, Gu Q, Sze KH, Chu IK, Kao RYT, Lee KC, Lam CW, Yang D, Shing-Chiu Tai S, Ke Y, Chan E, Chan WM, Dai J, Leung SP, Leung SY, Yuen KY. 2017. A tricyclic pyrrolobenzodiazepine produced by *Klebsiella oxytoca* is associated with cytotoxicity in antibiotic-associated hemorrhagic colitis. Journal of Biological Chemistry 292:19503–19520.

11. Martınez-Antonio A, Collado-Vides J. 2003. Identifying global regulators in transcriptional regulatory networks in bacteria. Curr Opin Microbiol 6:482–489.

12. Ishihama A, Shimada T, Yamazaki Y. 2016. Transcription profile of *Escherichia coli*: genomic SELEX search for regulatory targets of transcription factors. Nucleic Acids Res 44:2058–2074.

13. Unoarumhi Y, Blumenthal RM, Matson JS. 2016. Evolution of a global regulator: Lrp in four orders of γ-Proteobacteria. BMC Evol Biol 16:1–12.

14. Rodríguez-Valverde D, León-Montes N, Soria-Bustos J, Martínez-Cruz J, González-Ugalde R, Rivera-Gutiérrez S, González-Y-Merchand JA, Rosales-Reyes R, García-Morales L, Hirakawa H, Fox JG, Girón JA, De la Cruz MA, Ares MA. 2021. cAMP Receptor Protein Positively Regulates the Expression of Genes Involved in the Biosynthesis of *Klebsiella oxytoca* Tilivalline Cytotoxin. Front Microbiol 12:743594.

15. Seshasayee ASN, Sivaraman K, Luscombe NM. 2011. An overview of prokaryotic transcription factors. A handbook of transcription factors 7–23.

16. Kroner GM, Wolfe MB, Freddolino PL. 2019. *Escherichia coli* Lrp Regulates One-Third of the Genome via Direct, Cooperative, and Indirect Routes. J Bacteriol 201.

17. Kutukova EA, Livshits VA, Altman IP, Ptitsyn LR, Zyiatdinov MH, Tokmakova IL, Zakataeva NP. 2005. The yeaS (leuE) gene of *Escherichia coli* encodes an exporter of leucine, and the Lrp protein regulates its expression. FEBS Lett 579:4629–4634.

18. Lintner RE, Mishra PK, Srivastava P, Martinez-Vaz BM, Khodursky AB, Blumenthal RM. 2008. Limited functional conservation of a global regulator among related bacterial genera: Lrp in *Escherichia, Proteus and Vibrio*. BMC Microbiol 8:1–26.

19. Hart BR, Blumenthal RM. 2011. Unexpected Coregulator Range for the Global Regulator Lrp of *Escherichia coli* and *Proteus mirabilis*. J Bacteriol 193:1054–1064.

20. Cho BK, Barrett CL, Knight EM, Park YS, Palsson B. 2008. Genome-scale reconstruction of the Lrp regulatory network in Escherichia coli. Proc Natl Acad Sci U S A 105:19462–19467.

21. Chen S, Calvo JM. 2002. Leucine-induced dissociation of *Escherichia coli* Lrp hexadecamers to octamers. J Mol Biol 318:1031–1042.

22. Lee E, Pokoo R, Nguyen LT, Lee CY. 2017. Analysis of quaternary structure of leucine-responsive regulatory protein (Lrp) by crosslink experiments. Korean Journal of Microbiology 53:297–303.

23. Berthiaume F, Crost C, Labrie V, Martin C, Newman EB, Harel J. 2004. Influence of L -Leucine and L-Alanine on Lrp Regulation of *foo*, Coding for F165, a Pap Homologue. J Bacteriol 186:8537–8541.

24. McFarland KA, Lucchini S, Hinton JCD, Dorman CJ. 2008. The leucine-responsive regulatory protein, Lrp, activates transcription of the fim operon in *Salmonella enterica* serovar Typhimurium via the *fimZ* regulatory gene. J Bacteriol 190:602– 612.

25. Tani TH, Khodursky A, Blumenthal RM, Brown PO, Matthews RG. 2002. Adaptation to famine: a family of stationary-phase genes revealed by microarray analysis. Proceedings of the National Academy of Sciences 99:13471–13476.

26. Huang I, Chen K-Y, Rathod J, Chiu Y-C, Chen J-W, Tsai P-J. 2019. The transcriptional regulator Lrp contributes to toxin expression, sporulation, and swimming motility in *Clostridium difficile*. Front Cell Infect Microbiol 9:356.

27. Darby A, Lertpiriyapong K, Sarkar U, Seneviratne U, Park DS, Gamazon ER, Batchelder C, Cheung C, Buckley EM, Taylor NS, Shen Z, Tannenbaum SR, Wishnok JS, Fox JG. 2014. Cytotoxic and pathogenic properties of *Klebsiella oxytoca* isolated from laboratory animals. PLoS One 9.

28. Perez JC, Groisman EA. 2007. Acid pH activation of the PmrA/PmrB two-component regulatory system of *Salmonella enterica*. Mol Microbiol 63:283–293.

29. Chen S, Iannolo M, Calvo JM. 2005. Cooperative binding of the leucine-responsive regulatory protein (Lrp) to DNA. J Mol Biol 345:251–64.

30. Madeira F, Pearce M, Tivey ARN, Basutkar P, Lee J, Edbali O, Madhusoodanan N, Kolesnikov A, Lopez R. 2022. Search and sequence analysis tools services from EMBL-EBI in 2022. Nucleic Acids Res 10.1093/nar/gkac240.

31. Datsenko KA, Wanner BL. 2000. One-step inactivation of chromosomal genes in *Escherichia coli* K-12 using PCR products. Proc Natl Acad Sci U S A 97:6640– 6645.

32. Jahn CE, Charkowski AO, Willis DK. 2008. Evaluation of isolation methods and RNA integrity for bacterial RNA quantitation. J Microbiol Methods 75:318–324.

33. Aranda PS, LaJoie DM, Jorcyk CL. 2012. Bleach gel: A simple agarose gel for analyzing RNA quality. Electrophoresis 33:366–369.

34. Livak KJ, Schmittgen TD. 2001. Analysis of relative gene expression data using real-time quantitative PCR and the 2− ΔΔCT method. methods 25:402–408.

35. Schmittgen TD, Livak KJ. 2008. Analyzing real-time PCR data by the comparative C T method. Nat Protoc 3:1101.

36. Chimal-Cázares F, Hernández-Martínez G, Pacheco S, Ares MA, Soria-Bustos J, Sánchez-Gutiérrez M, Izquierdo-Vega JA, Ibarra JA, González-Y-Merchand JA, Gorvel J-P, Méresse S, De la Cruz MA. 2020. Molecular Characterization of SehB, a Type II Antitoxin of *Salmonella enterica* Serotype Typhimurium: Amino Acid Residues Involved in DNA-Binding, Homodimerization, Toxin Interaction, and Virulence. Front Microbiol 11:614.

37. Torres Montaguth OE, Bervoets I, Peeters E, Charlier D. 2019. Competitive Repression of the artPIQM Operon for Arginine and Ornithine Transport by Arginine Repressor and Leucine-Responsive Regulatory Protein in *Escherichia coli*. Front Microbiol 10:1563.

38. Brinkman AB, Ettema TJG, de Vos WM, van der Oost J. 2003. The Lrp family of transcriptional regulators. Mol Microbiol 48:287–94.

39. Ambartsoumian G, D’Ari R, Lin RT, Newman EB. 1994. Altered amino acid metabolism in lrp mutants of *Escherichia coli* K12 and their derivatives. Microbiology (Reading) 140 ( Pt 7):1737–44.

40. Newman EB, Lin R. 1995. Leucine-responsive regulatory protein: a global regulator of gene expression in *E. coli*. Annu Rev Microbiol 49:747–75.

41. Beloin C, Ayora S, Exley R, Hirschbein L, Ogasawara N, Kasahara Y, Alonso JC, Hégarat FL. 1997. Characterization of an *lrp*-like (*lrpC*) gene from *Bacillus subtilis*. Mol Gen Genet 256:63–71.

42. Chen KY, Rathod J, Chiu YC, Chen JW, Tsai PJ, Huang IH. 2019. The Transcriptional Regulator Lrp Contributes to Toxin Expression, Sporulation, and Swimming Motility in *Clostridium difficile*. Front Cell Infect Microbiol 9:356.

43. Kaiser JC, Heinrichs DE. 2018. Branching Out: Alterations in Bacterial Physiology and Virulence Due to Branched-Chain Amino Acid Deprivation. mBio 9.

44. Amorim Franco TM, Blanchard JS. 2017. Bacterial Branched-Chain Amino Acid Biosynthesis: Structures, Mechanisms, and Drugability. Biochemistry 56:5849– 5865.

45. Liu Y-K, Kuo H-C, Lai C-H, Chou C-C. 2020. Single amino acid utilization for bacterial categorization. Sci Rep 10:12686.

46. Friedberg D, Platko J v., Tyler B, Calvo JM. 1995. The amino acid sequence of Lrp is highly conserved in four enteric microorganisms. J Bacteriol 177:1624–1626.

47. Ishii Y, Shige Y, Akasaka N, Trinugraha AC, Higashikubo H, Fukuda W, Fujiwara S. 2021. Leucine-Responsive Regulatory Protein in Acetic Acid Bacteria Is Stable and Functions at a Wide Range of Intracellular pH Levels. J Bacteriol 203.

48. Ares MA, Fernández-Vázquez JL, Pacheco S, Martínez-Santos VI, Jarillo-Quijada MD, Torres J, Alcántar-Curiel MD, González-Y-Merchand JA, De La Cruz MA. 2017. Additional regulatory activities of MrkH for the transcriptional expression of the *Klebsiella pneumoniae mrk* genes: Antagonist of H-NS and repressor. PLoS One 12.

49. Yokoyama K, Ishijima SA, Clowney L, Koike H, Aramaki H, Tanaka C, Makino K, Suzuki M. 2006. Feast/famine regulatory proteins (FFRPs): *Escherichia coli* Lrp, AsnC and related archaeal transcription factors. FEMS Microbiol Rev 30:89–108.

50. Calvo JM, Matthews RG. 1994. The leucine-responsive regulatory protein, a global regulator of metabolism in *Escherichia coli*. Microbiol Rev 58:466–490.

51. Ziegler CA, Freddolino PL. 2021. The leucine-responsive regulatory proteins/feast-famine regulatory proteins: an ancient and complex class of transcriptional regulators in bacteria and archaea. Crit Rev Biochem Mol Biol 56:373–400.

52. Baek C-H, Wang S, Roland KL, Curtiss R. 2009. Leucine-Responsive Regulatory Protein (Lrp) Acts as a Virulence Repressor in *Salmonella enterica* Serovar Typhimurium. J Bacteriol 191:1278–1292.

53. Minami J, Okabe A, Shiode J, Hayashi H. 1989. Production of a unique cytotoxin by *Klebsiella oxytoca*. Microb Pathog 7:203–211.

54. Higaki M, Chida T, Takano H, Nakaya R. 1990. Cytotoxic component(s) of *Klebsiella oxytoca* on HEp-2 cells. Microbiol Immunol 34:147–151.

55. Beaugerie L, Metz M, Barbut F, Bellaiche G, Bouhnik Y, Raskine L, Nicolas JC, Chatelet FP, Lehn N, Petit JC. 2003. *Klebsiella oxytoca* as an agent of antibiotic-associated hemorrhagic colitis. Clinical Gastroenterology and Hepatology 1:370– 376.

56. Cowles KN, Goodrich-Blair H. 2005. Expression and activity of a *Xenorhabdus nematophil*a haemolysin required for full virulence towards Manduca sexta insects. Cell Microbiol 7:209–219.

57. Casadaban MJ. 1976. Transposition and fusion of the lac genes to selected promoters in *Escherichia coli* using bacteriophage lambda and Mu. J Mol Biol 104:541–555.

58. Mayer MP. 1995. A new set of useful cloning and expression vectors derived from pBlueScript. Gene 163:41–46.

